# RNASeq as a tool to understand dysregulation of potential biomarkers in HNSC

**DOI:** 10.1101/2022.09.20.508683

**Authors:** Ezhuthachan Mithu, Madhvi Joshi, Ishan Raval, Chaitanya Joshi, Anirban Dasgupta, Sharmistha Majumdar, Siddharth A Shah

**Affiliations:** Gujarat Biotechnology Research Centre, MS Building, Gandhinagar 382010, India; Indian Institute of Technology, Gandhinagar; Zydus Cancer Centre, Thaltej, Ahmedabad, Gujarat

**Author notes:** **Correspondence:** Dr. Madhvi Joshi.

**Keywords:** Oral Squamous cell carcinoma, HNSC (Head and Neck Squamous Cell Carcinoma), Biomarker, Margin clearance, Hub genes, TCGA(The Cancer Genome Atlas)

## Abstract

With its rising fatality rates, oral cancer is one of the most concerning public health issues. To reduce disease-related mortality and morbidity, advancements in screening and detection are critical. Finding specific biomarkers is one of the most successful approaches for screening, diagnosing, and staging this dreadful disease. In this study differentially expressed genes associated with oral cancer were analyzed using RNASeq to find the potential biomarkers. Functional enrichment of upregulated genes found that 253 genes were present in the plasma membrane. Three clusters were formed using KMean Clustering from the PPI networks, and highly connected hub genes were identified from each cluster. Eventually, expression and survival analyses of hub genes were performed using The Cancer Genome Atlas (TCGA) database targeting Head and Neck Squamous Cell Carcinoma. Among those genes, expression levels of eight genes *SLC2A1, ITGA6, LAMC2, COL1A2, COL1A1, TNC, THY1*, and *CD276* have significantly changed in Head and Neck Squamous cell carcinoma. There are reports that suggest these genes were significantly dysregulated in Oral Squamous cell carcinoma and can be explored further as potential biomarkers for margin clearance.

## 1 Introduction

Oral cancer is a malignant neoplasm affecting the lip, the floor of the mouth, cheek, gingiva, palate, or tongue (1). As per the report by Jemal et al., 2011 (2), nearly 2,63,900 new cases and 1,28,000 deaths occur due to oral cancer. Oral cancer is caused by smoking, alcohol drinking, tobacco chewing, and poor oral hygiene. In developing countries like India, oral cancer is a big concern as 20 per 100000 populations are affected by Oral cancer i.e. nearly 30% of all types of cancer [1]. India is the country with the second-highest rate of mouth cancer. Mostly this cancer is diagnosed at later stages, resulting in low treatment outcomes and also high costs. As a result, oral cancer has a dismal prognosis when diagnosed late. Early detection and knowledge dissemination are essential in the diagnosis of oral cancer (3). To date, the standard diagnostic procedures used for diagnosing oral malignancy and lesions are scalpel biopsy and histopathological examination (4). This traditional method is invasive making patients discomfort along with a high recurrence rate (5). In the past few decades use of biomarkers in diagnosing and treatment is increasing widely. The surface markers can be used as diagnostic markers as well as in drug targets(6). Biomarkers can be grouped into three classes prognostic markers, predictive markers, and theranostic markers. A prognostic marker is used to understand disease occurrence and progression, a predictive marker is used for the early assessment of potential therapy and a theranostic marker combines both prognostic and predictive markers [6]. Considering total human proteins approximately 30 % of proteins are membrane proteins, but only a few proteins are used as biomarkers for the diagnosis of cancers. Membrane proteins are classified based on cellular localization, protein interactions, motifs, cellular functions, or interactions. Membrane proteins have the potential of representing all these biomarkers, thus gaining increasing medical diagnostic applications (7). In oral cancer, in vivo research or single-gene studies are highly preferred, with an impact on the whole transcriptome being rare. The present study is primarily focused on identifying highly elevated genes on the plasma membrane and assessing differentially expressed genes in tumor tissue and adjacent normal tissue. PPI network and MCode analysis of these genes were utilized to uncover genes from three clusters, and those genes were validated using the TCGA (The Cancer Genome Atlas) database of Head and Neck Squamous cell carcinoma.

## 2 Materials and Methods

### 2.1 Collection of Tissues

Four pairs of Oral Squamous cell carcinoma tissues and normal adjacent tissues were collected from Zydus Cancer Centre in Ahmedabad, Gujarat, India, with written informed consent from all patients. Before being used, all tissue specimens were kept in RNAlater vials and then stored at −80°C until further use.

### 2.2 Total RNA extraction

The total RNA from each tissue was extracted by RNeasy Mini kit (Qiagen) as per the manufacturer’s instruction (8). The concentration of the extracted RNA was evaluated by Qubit (Invitrogen, USA) and the RNA integrity was checked by Bioanalyzer (Agilent 2100). To avoid contamination, RNA extraction was carried out utilizing established procedures.

### 2.3 RNA Sequencing

For RNA sequencing, four pairs of tumor and adjacent normal tissues were obtained. The Ribosomal RNA (rRNA) from total RNA was removed after RNA extraction using Low Input RiboMinus^™^ Eukaryote System v2, and the library was generated using the Illumina TruSeq Stranded Total RNA kit (Illumina, CA, USA) as directed by the manufacturer (9). To synthesize the initial strand of cDNA, the RNA was fragmented and then primed with random hexamers. The second strand was then synthesized, followed by the production of blunt-ended cDNA. Following that, the 3’end of the dsDNA fragment was adenylated to prevent the adaptor ligation process from ligating with each other. Multiple indexing adapters were subsequently ligated to the ends of the ds cDNA segments. The ligated fragments were cleaned using AMPure XP beads before being enriched with a thermal cycler to increase the amount of DNA in the library. The fragments were sequenced by Illumina MiSeq (Illumina, CA, USA) using MiSeq V2 300 cycle reagent kit after the amplified library was purified with AMPure XP beads (Beckman Coulter, CA, USA) (Illumina, CA, USA) (10).

### 2.4 RNA-Sequencing data processing and analysis

Following Illumina MiSeq Next-Generation Sequencing, the quality of the acquired reads was evaluated using FastQC tool, and the data were then trimmed using PrinSeq followed by quality check using FastQC. The sequenced reads were mapped with the hg38 human genome and sequence analysis was done by the CLC genomics workbench (12.0.3, Qiagen Bioinformatics). Additionally, DEseq2 in RStudio was used to conduct a differential gene expression between the Patient’s normal adjacent tissue and a healthy volunteer’s oral squamous tissue (11).

### 2.5 GO Enrichment analysis, GO annotation, and functional enrichment

Cluster Profiler in R was used to perform GO enrichment of all significantly expressed genes on the basis of log2fold change and padj value 0.05 (12). WebGeStalt was used to annotate the genes that were significantly upregulated and downregulated (13). FunRich 3.1.3 (software programme for functional enrichment of genes), was used for the functional enrichment of genes (14).

### 2.6 Protein−Protein Interaction (PPI) Network Construction

Using the Search Tool for the Retrieval of Interacting Genes (STRING, version 11.5), the upregulated genes discovered in Plasma Membrane were used to create PPI networks (15). Based on the distance matrix, three clusters were created, each including genes with high global scores.

### 2.7 Identification of Hub genes

From three different clusters obtained, the hub genes were identified by the Molecular Complex Detection (MCODE) plugin for three different clusters (16).

### 2.8 Validation of hub genes

Gene Expression Profiling Interactive Analysis (GEPIA) allows for a comparison of the survival effect and DEG expression profile analysis in oral cancer (17). To validate the aforementioned hub genes, the relative expression levels in head and neck squamous cell carcinoma (HNSC) was done with cutoff of |log2FC| > 2 and a p-value of 0.05. Furthermore, the overall survival (OS) effect of hub genes in HNSC was calculated using the log-rank p-value and hazard ratio (HR). To split patients into high (more than the 75th percentile) and low (greater than the 25th percentile) expression groups, the normalized expression level of selected genes across all HNSC patients in TCGA database was collected. Since, both GEPIA and TCGA database do not includes OSCC category, HNSC was used for the validation studies, and OSCC accounts for 95% of HNSC (18).

## 3 Results

### 3.1 Characteristics of Study Subjects

This study included 4 study subjects, from which subjects were of different stages. Tumor tissue and adjacent normal tissue was collected from each individual by the surgeon (Table1).

**Table 1.** The characteristics of the study subjects including harmful past habits leading to oral cancer.

|  | Sex | Harmful Habits | Cancer Stage | Tissue type |
| --- | --- | --- | --- | --- |
| SRR19894454 | Male | Tobacco Chewing | Stage IV | Adjacent Normal Tissue |
| SRR19894453 | Male | Tobacco Chewing | Stage IV | Tumor Tissue |
| SRR19894450 | Male | Tobacco Chewing | Stage III | Adjacent Normal Tissue |
| SRR19894449 | Male | Tobacco Chewing | Stage III | Tumor Tissue |
| SRR19894448 | Male | Tobacco Chewing | Stage II | Adjacent Normal Tissue |
| SRR19894447 | Male | Tobacco Chewing | Stage II | Tumor Tissue |
| SRR19894452 | Male | Tobacco Chewing | Stage IV | Adjacent Normal Tissue |
| SRR19894451 | Male | Tobacco Chewing | Stage IV | Tumor Tissue |

### 3.2 RNA Quality and Sequencing

The extracted RNA of 8 Samples had RIN Value between 7 to 9. After Library preparation and sequencing the average base pair obtained was around 350 bp. Percentage of good quality sequences after filtration was nearly 90 to 98% with around 50-55 GC %.

### 3.3 Identification of Differentially expressed genes

DEGs (Differentially expressed genes) were discovered using characteristics such as absolute log2foldchange >2 and padj value< 0.05, yielding 1518 upregulated genes and 2226 downregulated genes Figure 1.A. As a result, a heat map was created using RPKM values, as shown in Figure 1.B. GO enrichment of relevant genes was performed using cluster profile in R, and the analysis revealed that the majority of genes were suppressed in supramolecular complex, cytoskeletal protein binding, and system process. While several genes were identified to be activated in processes like androgen catabolism, interleukin-1 receptor binding, and CXCR3 chemokine receptor binding (Figure2). GO enrichment of relevant genes was performed using cluster profile in R, and the analysis revealed that the majority of genes were suppressed in supramolecular complex, cytoskeletal protein binding, and system process. While several genes were identified to be activated in processes like androgen catabolism, interleukin-1 receptor binding, and CXCR3 chemokine receptor binding (Figure2).

**Figure 1.**
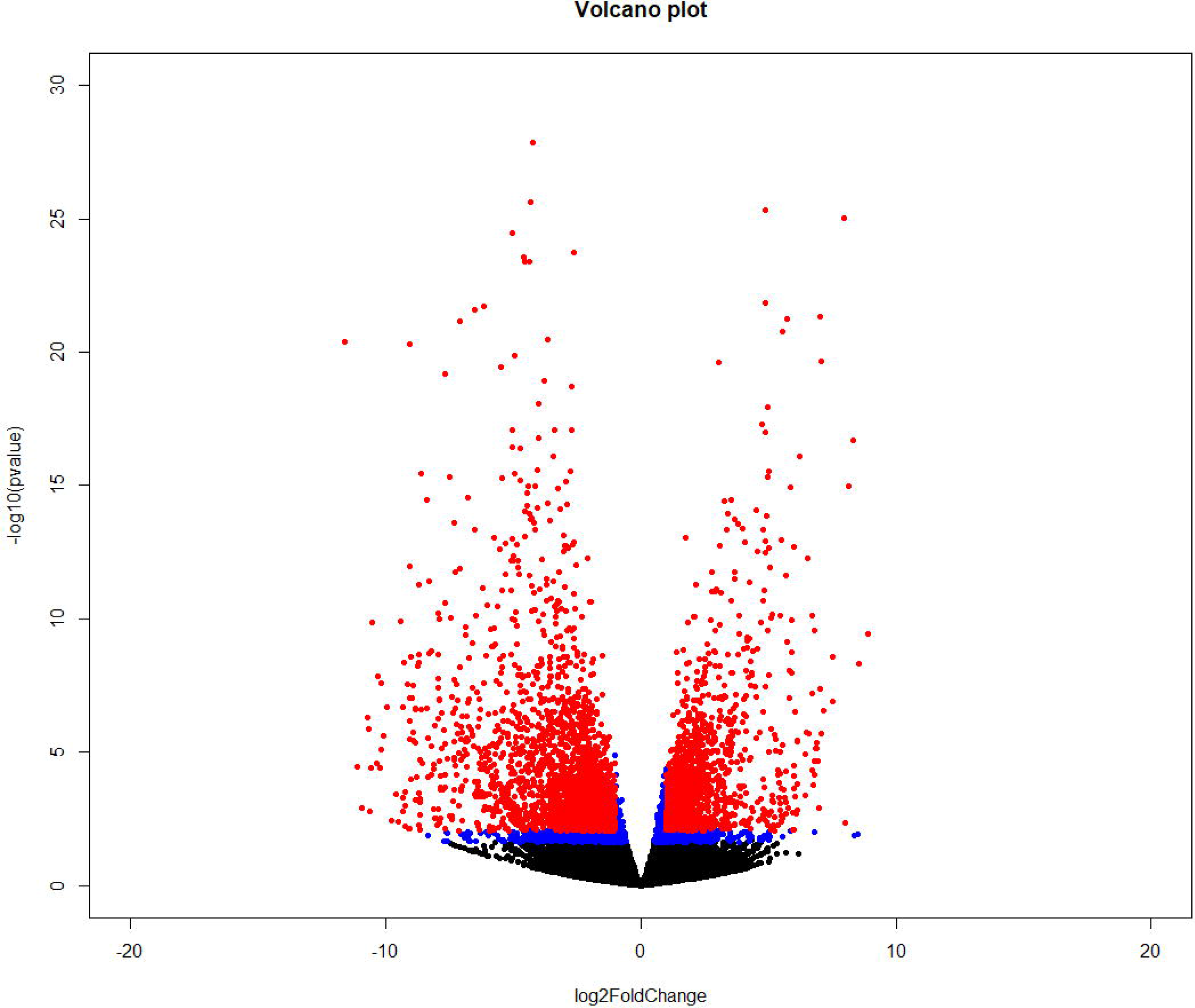
Identification of differentially expressed genes (A) Volcano plot generated from DeSeq2 analysis of four Patient samples having Tumor tissue and Adjacent normal tissue. (B) Heat Map generated from the RPKM values of Differentially expressed genes obtained when Tumor tissue was compared with normal tissue.

**Figure 2.**
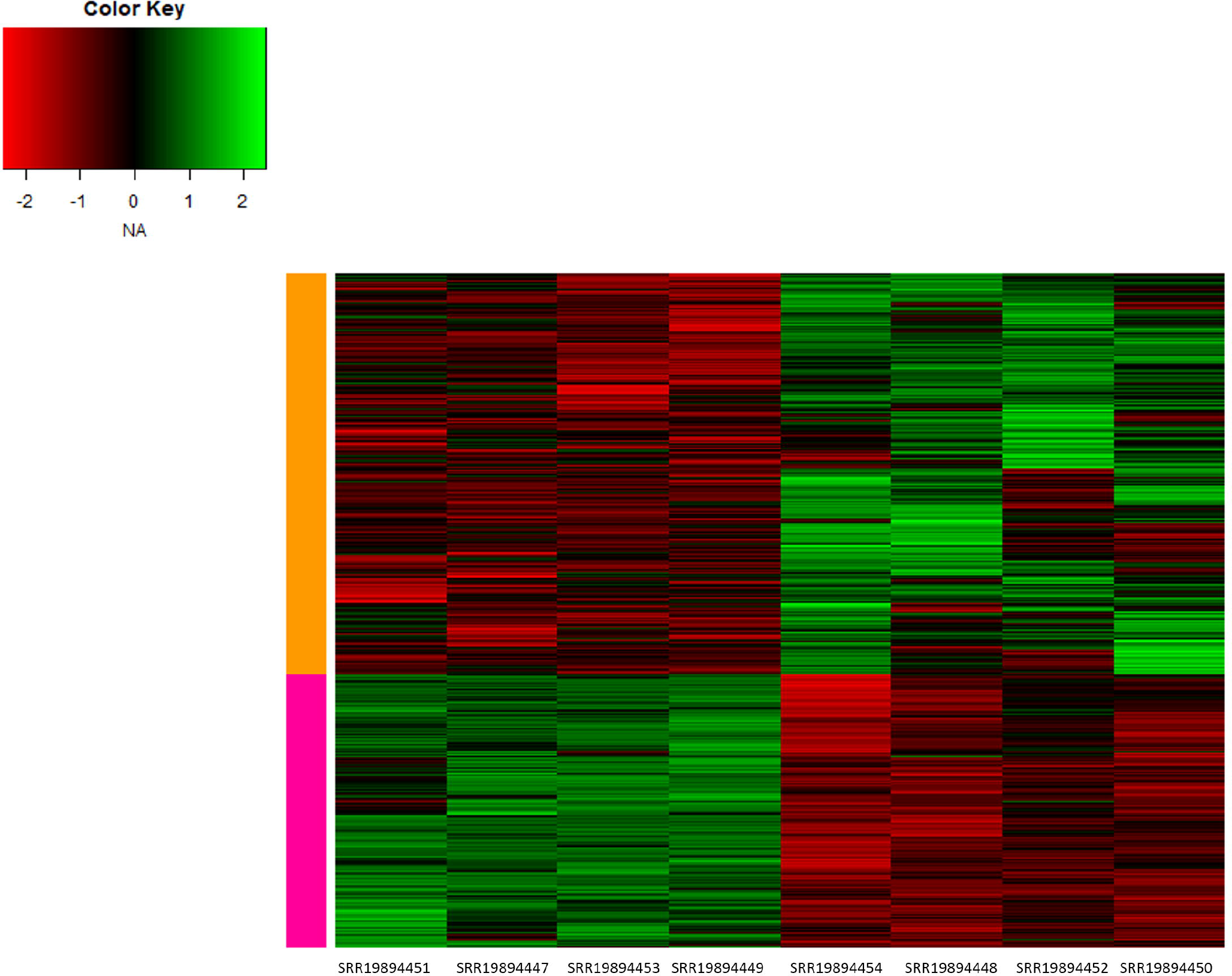
GO enrichment analysis for all the significant genes which are dysregulated using ClusterProfiler in R

### 3.4 GO annotation and Functional Enrichment

WebGeStalt was utilized to annotate the genes that were significantly upregulated and downregulated. A total of 1304 genes were classified in the category of upregulated genes. 827 genes were annotated for biological regulation and 817 genes for metabolic process in biological process. From the identified genes in cellular components, 602 genes were located in the membrane and 560 genes in the nucleus. The majority of genes were located in the Protein Binding domain, based on their molecular function (Figure 3.a). According to the GO annotation of downregulated genes, there were 1978 genes annotated, with 1147 genes annotated for biological regulation and 980 genes annotated for metabolic process, respectively. There were 887 genes detected in the membrane and 565 genes located in the nucleus among the annotated genes. 1031 genes were discovered to be involved in protein binding in molecular function (Figure 3.b).

**Figure 3.**
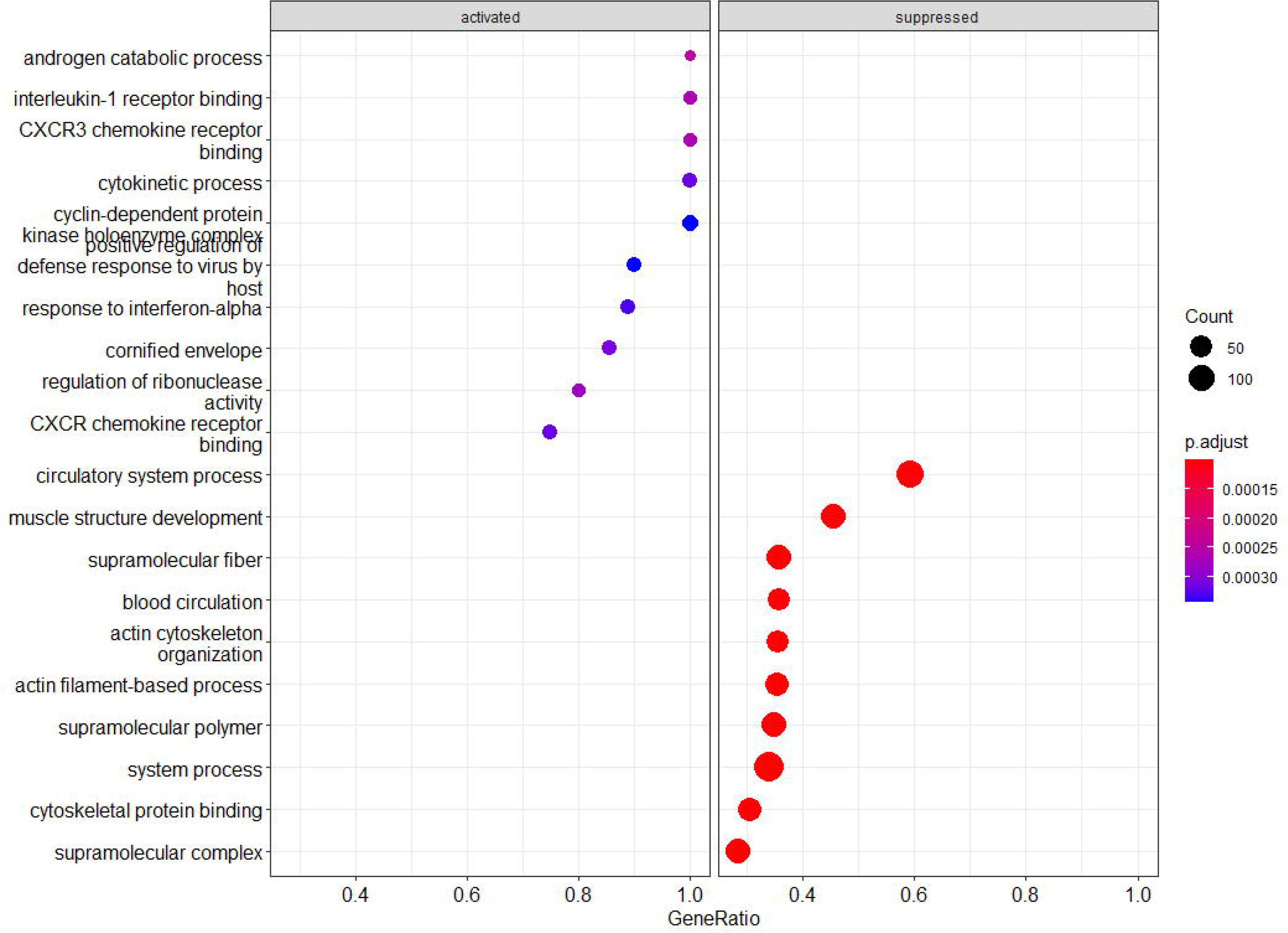
GO Annotation of Differentially expressed genes (A) GO annotation of Significant upregulated genes using WebGestalt. (B) GO annotation of significant downregulated genes using WebGestalt.

### 3.5 Functional Enrichment of Upregulated and Downregulated genes

Functional Enrichment of upregulated and downregulated genes is done using FunRich. The major goal was to look for elevated genes in plasma membranes and analyze them. From the analysis it was found that 253 genes were present in the plasma membrane which are significantly upregulated (Table 2).

**Table 2.**
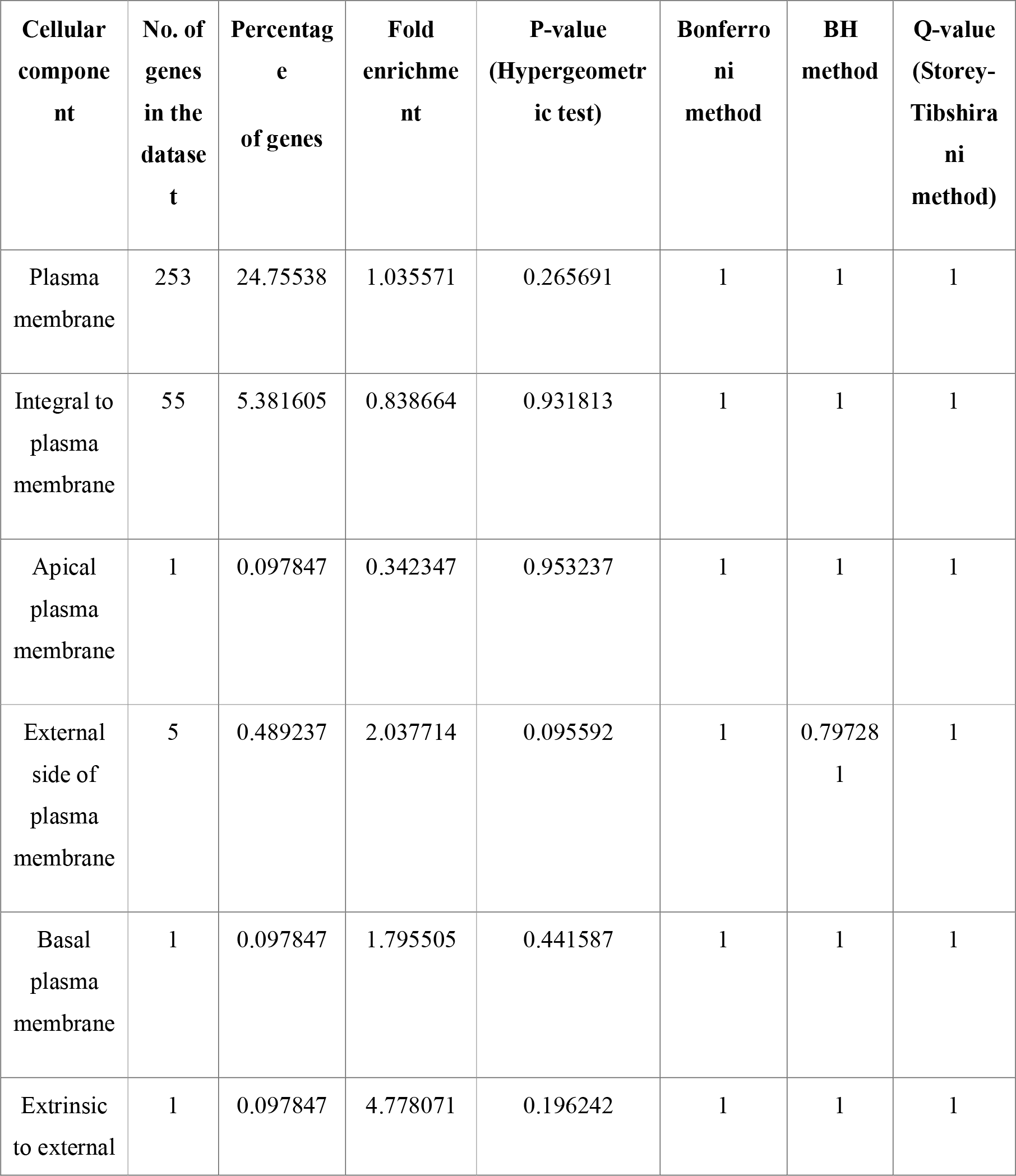
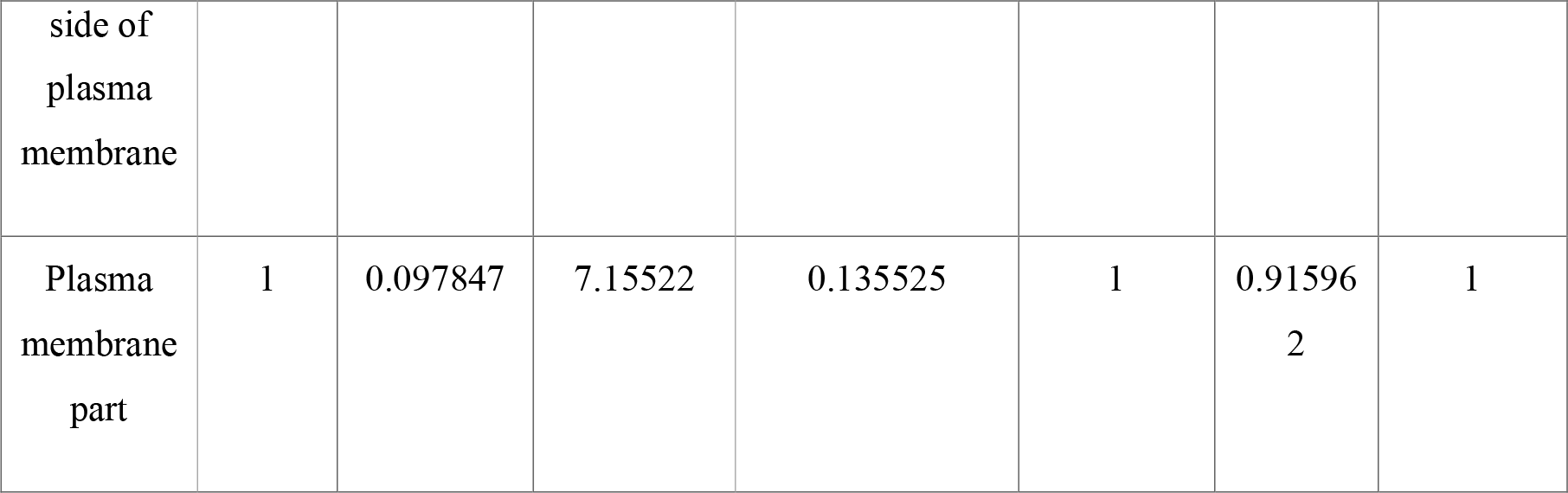
Functional enrichment of upregulated genes. The table describes the presence of 253 upregulated genes in Plasma membrane with fold enrichment value 1.035571.

### 3.6 Formation of PPI network and Identification of hub genes

The PPI network was built using the 253 genes found in the plasma membrane. Based on KMean clustering, the entire network was separated into three clusters. The set of genes is grouped into clusters based on their strongest similarity (Figure 4). There are primarily three clusters in this PPI network. Cytoscape’s Molecular Complex Detection (MCODE) plug-in was used analyze the three clusters. Nodes with a score of greater than 5.0 were chosen from three separate clusters. The blue cluster had 15 highly connected nodes with a score of 12.857 (Figure 5.a and Figure 5.b), the green cluster had 9 highly connected nodes with a score of 7.250 (Figure 6.a and Figure 6.b), and the red cluster had 20 highly interconnected nodes with a score of 17.474 (Figure 7.a and Figure 7.b).

**Figure 4.**
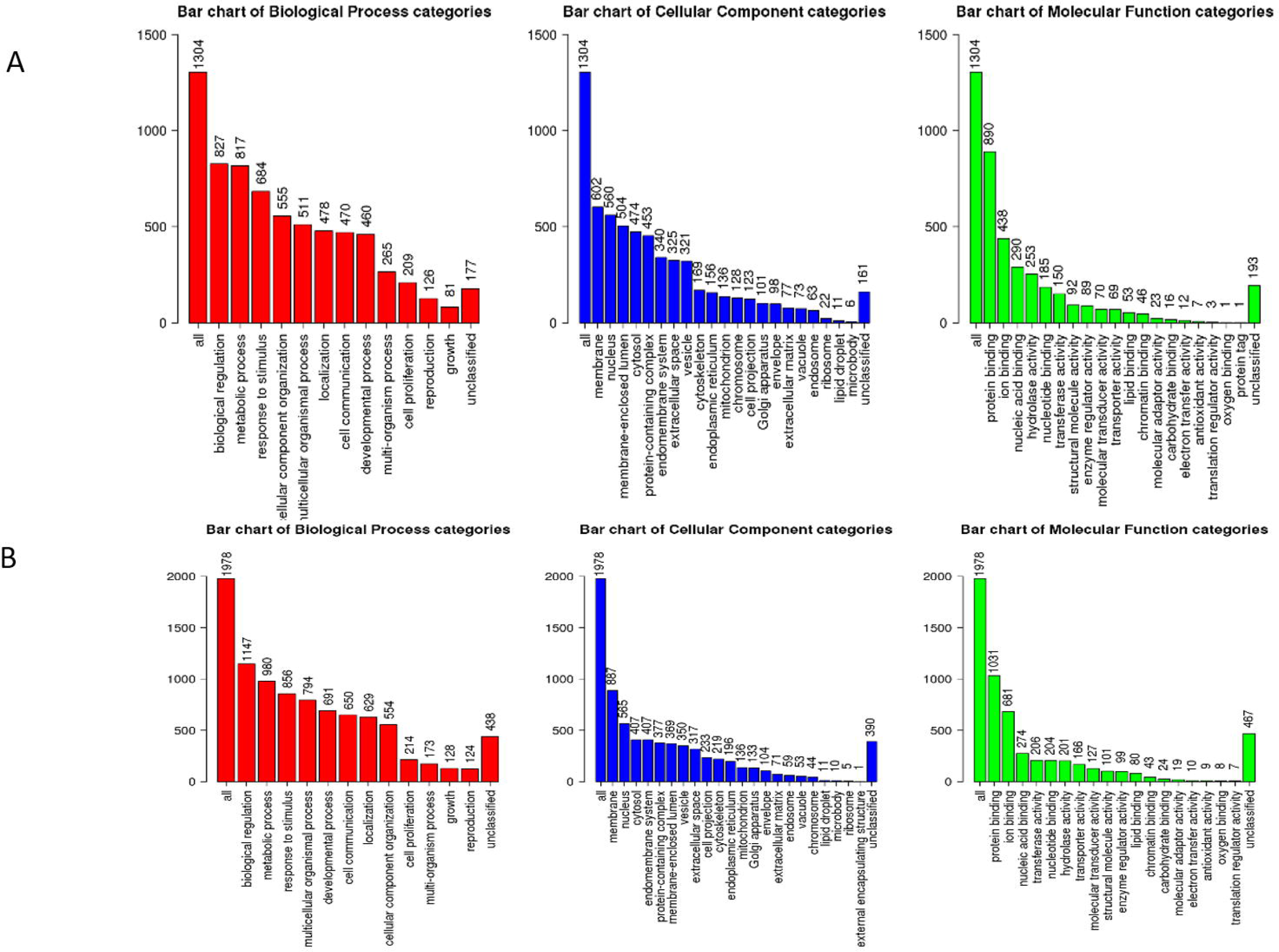
PPI Network of 253 genes analysed from the functional enrichment of upregulated genes in Plasma membrane

**Figure 5.**
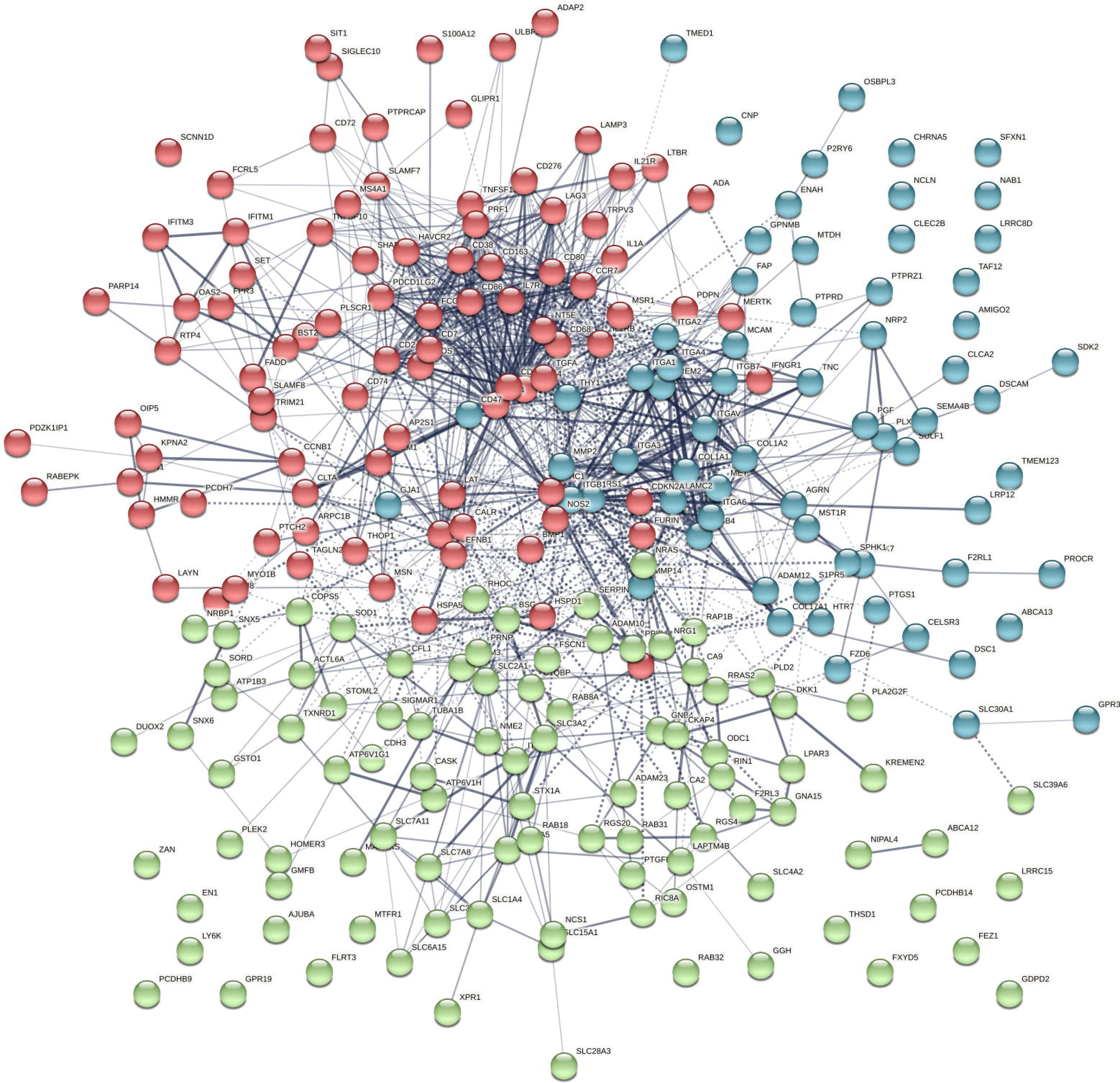
(A) Cluster 1 obtained from KMean clustering (B) MCode analysis of Cluster 1 found 15 nodes having score value 12.857

**Figure 6.**
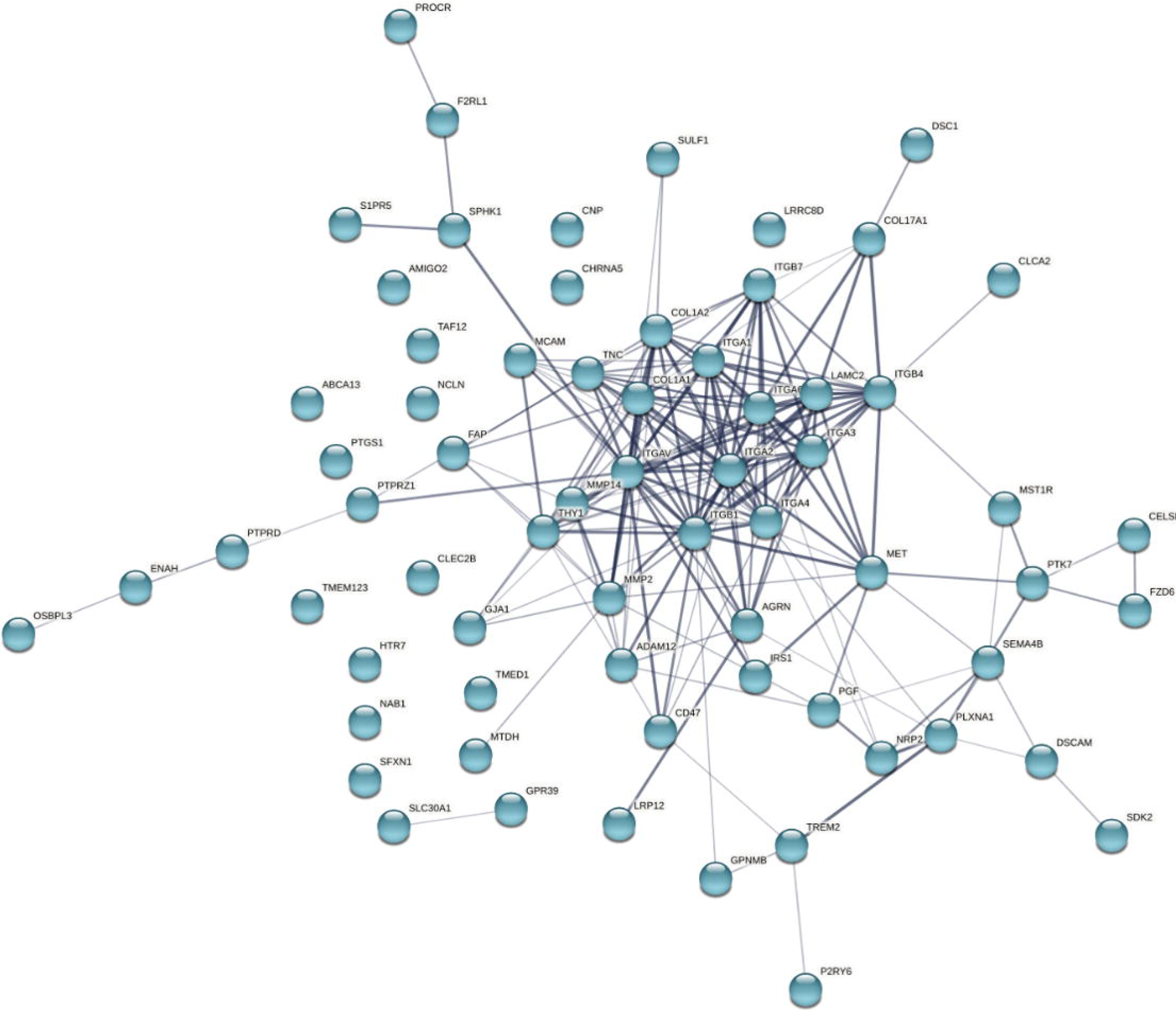
(A) Cluster 1 obtained from KMean clustering (B) MCode analysis of Cluster 1 found 9 nodes having score value 7.250

**Figure 7.**
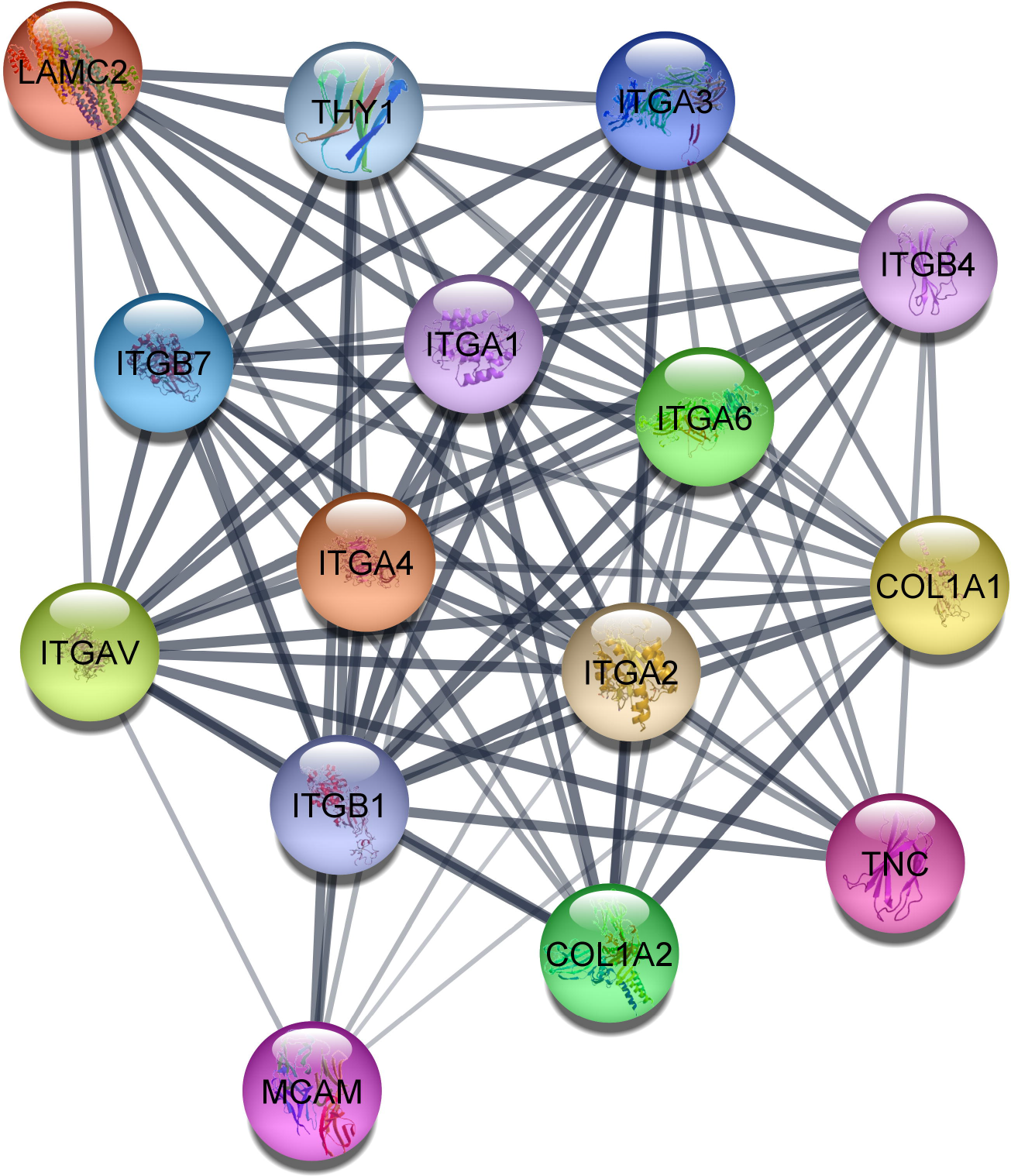
(A) Cluster 3 obtained from KMean clustering (B)MCode analysis of Cluster 3 found 20 nodes having score value 17.474

### 3.7 Validation of Hub genes

GEPIA2 was used to assess the clinical significance of genes discovered in three groups using The HNSC database. In the TCGA database, OSCC accounts for 95% of HNSC with log2foldchange>2 and padj value < 0.05. The expression level of genes found from three clusters were found to be significantly dysregulated in HNSC (Figure 8). Only those genes found to be highly significant from all the three clusters were used to determine prognostic value by comparing the levels of gene expression in tumour tissue and adjacent normal tissue. It was found that logrank P value of *SLC2A1, ITGA6, COL1A2, COL1A1, LAMC2, THY1, TNC* and *CD276* gene is 0.78, 0.37, 0.86, 0.3, 0.012, 0.59, 0.26 and 0.02 respectively. The HR-high (Hazard risk) *SLC2A1, ITGA6, COL1A2, COL1A1, LAMC2, THY1, TNC* and *CD276* majorly ranges between 1-1.5 (Figure 9).

**Figure 8.**
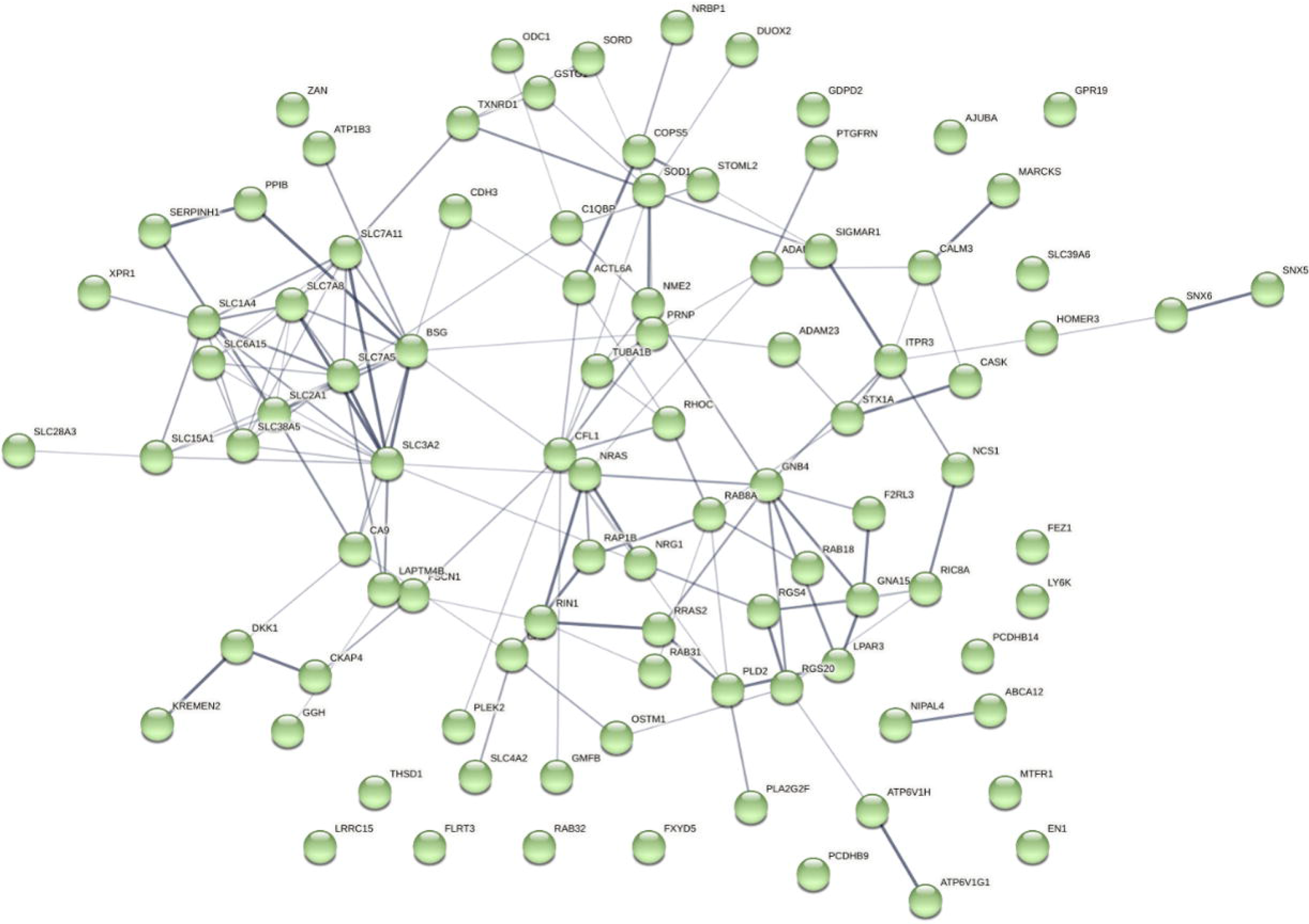
Gene expression of significant gene comparing Tumor tissue samples and normal tissue samples. The analysis was done using GEPIA2 targeting Head and Neck Squamous cell carcinoma in TCGA database

**Figure 9.**
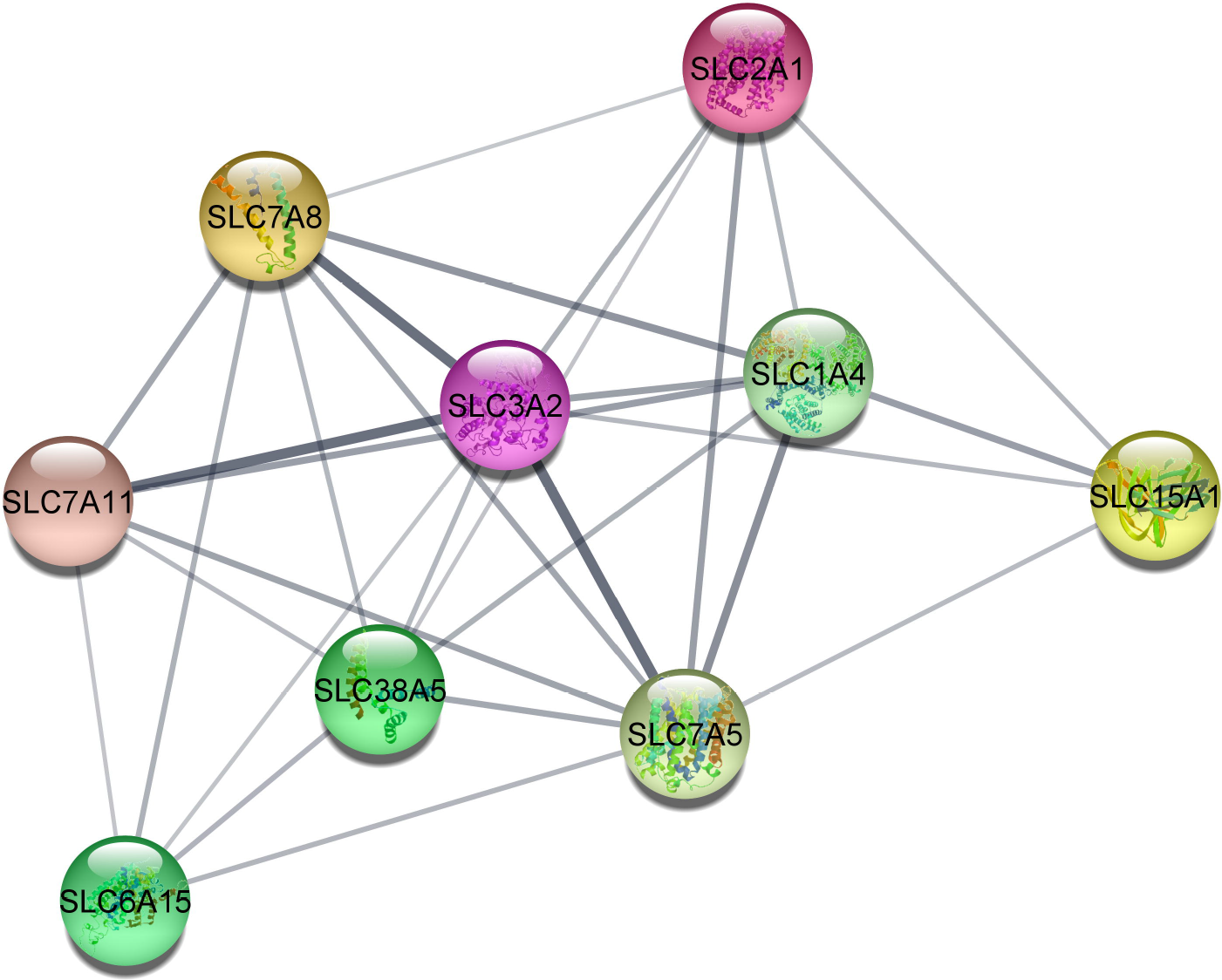
Kaplan-Meier survival analysis for high and low expression levels of *SLC2A1, ITGA6, LAMC2, COL1A2, COL1A1, TNC, THY1*, and *CD276* in Head and Neck Squamous Cell Carcinoma Patients from The Cancer Genome Atlas

## 4 Discussion

Oral cancer is ranked within the top 10 cancer on the global scale(19), prevailing more in men compared to women. Genomic instability is a key factor associated with oral cancer which leads to tumor progression [19]. Identification and characterization of genes involved in carcinogenesis will be a major hit in clinical management and drug treatment. The aim of this study was to determine whether the transcriptional expression of genes found in the plasma membrane could serve as biomarkers used for margin clearance. After sequencing, the reads mapped with hg38 were utilized to determine the dysregulated genes, which were then processed further. The study found that 1518 genes were upregulated and 2226 genes were downregulated. It was observed that with GO enrichment of dysregulated genes majority of genes from the gene sets were found to be activated in the androgen catabolic process. Androgen receptor (AR) functions as a ligand-dependent transcription factor, and AR binding to Androgen receptor elements activates genes involved in AR cell proliferation, differentiation, and survival. AR also promotes EGFR phosphorylation, which aids cell motility (20). GO enrichment also represents that genes are activated in Interleukin-1 receptor binding, IL1 is a pleiotropic cytokine that stimulates genes that promote cancer growth and metastasis(21).Functional enrichment of upregulated genes found that 253 genes significantly upregulated in the plasma membrane with a Fold Enrichment value of 1.035571 and a P-value of 0.265. *ADAM12, CA9, EN1, GDPD2, HTR7, MSR1, TNC, SLC2A1, ITGA6, LAMC2, COL1A2, COL1A1, CD276, TNC, THY1, CHN1*, and other genes were found to be most significant considering log2Foldchange >2 using GEPIA2 tool. There are reports suggesting the genes found to be significant by analysis using GEPIA2 tool found to be significantly associated with oral cancer. HER2/Erbb2 belong to member of epidermal growth factor receptor is resistance to chemotherapy and reduced survival of head and neck cancer. *ADAM12*(A Disintergrin and Metalloprotease12) is significantly correlated with *HER2*, and co-expression of *ADAM12* and *HER2* found in Head and Neck cancer(22). In OSCC (oral squamous cell carcinoma), by study it is reported that high *ITGA3* (integrin α3) was found in loco-regional dissemination and *ITGB4* (integrin β4) was highly expressed in distant metastasis(23). Using KMean clustering, the PPI network analysis of genes created three clusters. MCode analysis found the most correlated genes in all three clusters. When the GEPIA2 tool was used to validate genes from all clusters, it was discovered that they were all significantly dysregulated in tumor tissue compared to normal tissue.Cluster1 (Figure 5) represented *TNC, LAMC2, ITGA4, ITGB7, ITGA2, THY1, COL1A1, ITGA3, ITGB1, ITGAV, ITGA1, ITGA6, ITGB4, COL1A2, MCAM* are highly correlated. Among them *ITGA6, LAMC2, COL1A2, COL1A1, TNC, THY1* were the most significant genes. ECM (Extracellular matrix modelling) is a dynamic structure present in all tissues undergoing controlled remodeling (24). An uncontrolled remodelling of ECM leads to tumour growth and invasiveness (25). Integrin plays a role in ECM receptor interaction and focal adhesion, contributing to OSCC progression (26). *ITGA6* a member of integrin family protein functions as cell surface adhesion and signalling. Xie et al. 2021 (27) has reported that *ITGA6* (integrin subunit alpha 6) is overexpressed in Head and Neck Squamous Cell Carcinoma. Laminin, another gene found to be highly correlated in cluster1(figure 5) is a noncollagenous constituent of basement membrane, is involved in cell adhesion, signalling neurite outgrowth and metastasis. *LAMC2* is significantly associated with pattern and depth of invasion of squamous cell carcinoma. As per report of Nguyen et al. 2017 (28) *LAMC2* (laminin subunit gamma 2) was remarkably upregulated in Oral Squamous cell carcinoma at the cancer-stroma interface. Type 1 collagen is a key structural component of the extracellular matrix. Type 1 collagen is a heterotrimer of two type I α 1 (*COLIA1*) chains and one type I α 2 (*COLIA2*) chain. Abnormal expression of *COLIA1* and *COLIA2* is associated with Tumor progression. Lin et al. 2020 (29) reported that *COLIA1* and *COL1A2* is upregulated in HPSCC (hypopharyngeal squamous cell carcinoma) Cases. Increased deposition of *TNC* has been reported in tumor stroma of epithelial malignancies in mouth, colon, stomach and lung (30). Sundquist et al. 2017 (31) reported that in early stages of tumor patients *TNC* (tenascin-C) is negative, but with later stages *TNC* expression increases in Oral tongue squamous cell carcinoma. One of the most important stage in tumor metastasis is cancer cell adhesion to vascular endothelium. *Thy-1*(CD90) is a cell adhesion molecule which is activated in endothelial cells, and plays a vital role in melanoma metastasis by binding to integrins which are present in cancer cells (32).The related genes from Cluster 2 (figure 6) includes *SLC2A1, SLC7A8, SLC1A4, SLC15A1, SLC3A2, SLC38A5, SLC7A11, SLC6A15, SLC7A5*.The most significant gene *SLC2A1* (solute carrier family 2 member 1) known as *GLUT1* is crucial in cellular energy metabolic pathway. *SLC2A1* is over-expressed in several different types of carcinomas, including liver, lung, endometrial, oral and breast cancers, as well as gastric cancer (33).The related genes from Cluster3 (figure7) included *CD163, CD80, PDCD1LG2, CD38, CD4, CD276, CD274, NT5E, CD86, IL7R, CTLA4, IL2RB, HAVCR2, FCGR3A, CCR7, LAG3, ICOS, CD2, PRF1, CD68*. From Cluster3 (figure7) *CD276* is found to be the most significant. *CD276* molecule showed the highest significance in HNSC. *CD276* also known as the B7-H3 molecule, functions as an immune checkpoint belonging to the B7-CD28 pathway. The receptor CD276 is 110Kda, type 1 transmembrane glycoprotein having four Ig domains, with tandem duplication of IgV-IgC domain (4Ig-B7-H3) (34). Mainly, B7-H3 is expressed in immune cells, like Antigen-presenting Cells (APC). However, it is also present on Natural Killer cells, and B Cells (35). The reports suggest that it also acts as T-cell coinhibitory, inhibiting polyclonal or allogeneic CD4 and CD8 T-cell activation, proliferation, and effector cytokine production (36). The expression of *CD276* is upregulated upon IFNγ stimulation and Ig-linked transcript 4 signaling(37). The non-immunologic function of *CD276* regulates tumor aggressiveness. *CD276* modulates migration, invasion, and adhesion to fibronectin of various cancer cells by the Jak2/Stat3/MMP-9 signalling pathway (38). The expression of CD276 in Oral Squamous cell carcinoma tissue sample is 74.75% (39). As reported by Jung-Tsu et al., 2015 (40) N-gycans of *CD276* contains terminal α-galactose, having diverse fucosylation interacting better with DC-SIGN [DC-Specific intercellular adhesion molecule-3 (ICAM-3)-grabbing nonintegrin] and langerin on immune cells, enhancing the oral cancer. Yang et al., 2008 (41) also demonstrated that transfecting *CD276* plasmids into oral squamous cell carcinomas simultaneously enhances the proliferation of tumor cells, production of interferon, and the activity of cytotoxic T cells. These results give evidence of upregulation of *CD276* proliferating tumour and immune evasion serving as potential biomarkers for diagnosis of Oral Squamous cell carcinoma. We are cognizant that the limitation of our study includes in vitro validation of these genes using robust quantitative assays. However, taken together these findings establish a major highlight in classifying these significant genes promoting tumor cell progression and metastasis in the oral cavity and can be explored further as potential biomarkers for margin clearance.

## Author Contributions

ME conducted experiments, investigation, data curation, writing original draft. MJ and CJ contributed in conceptualization, funding, acquisition, supervision and project administration. SS did sample collection. IR, AD and SM did review and editing of manuscript. All authors contributed to the article and approved the submitted version.

---

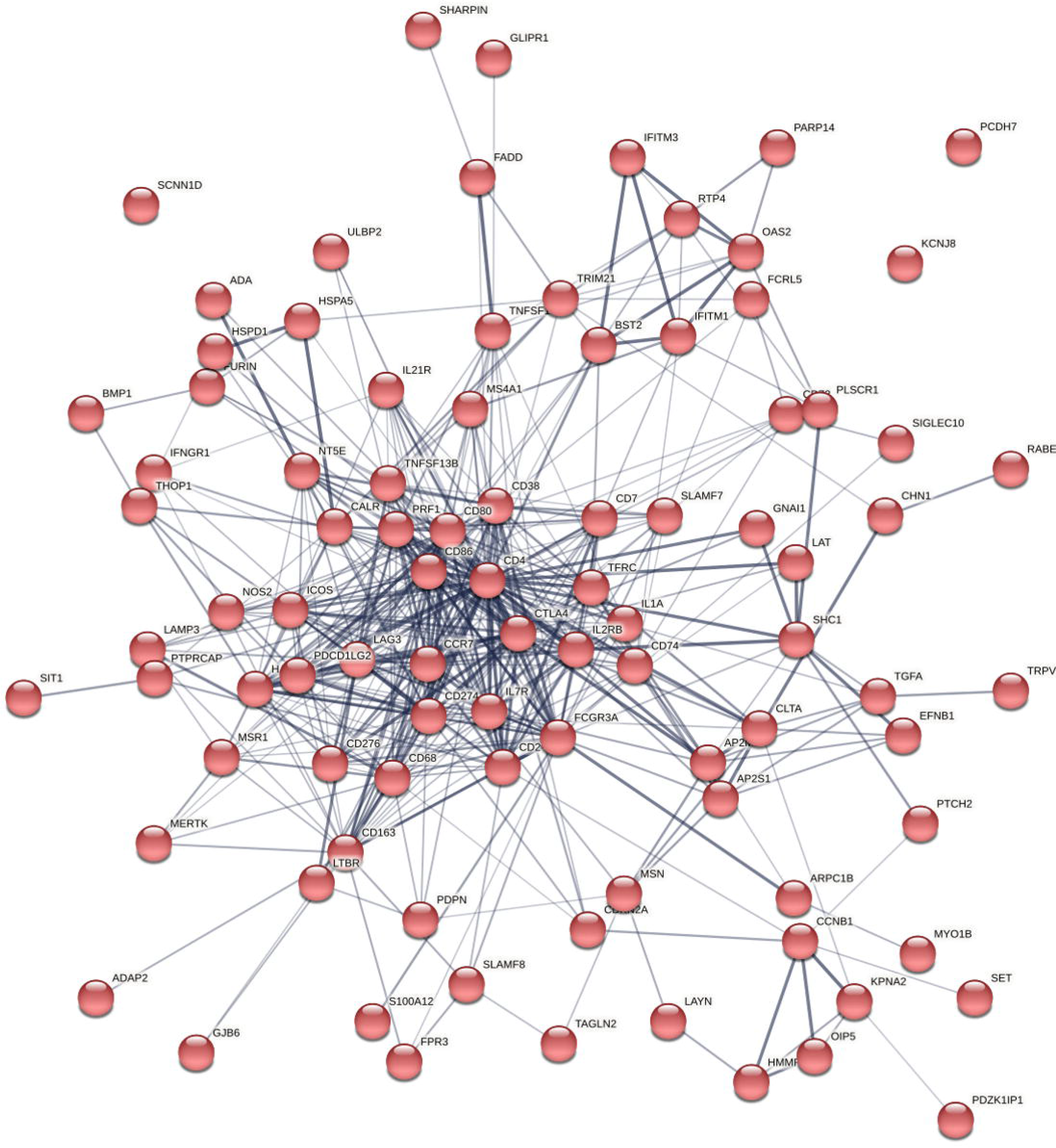

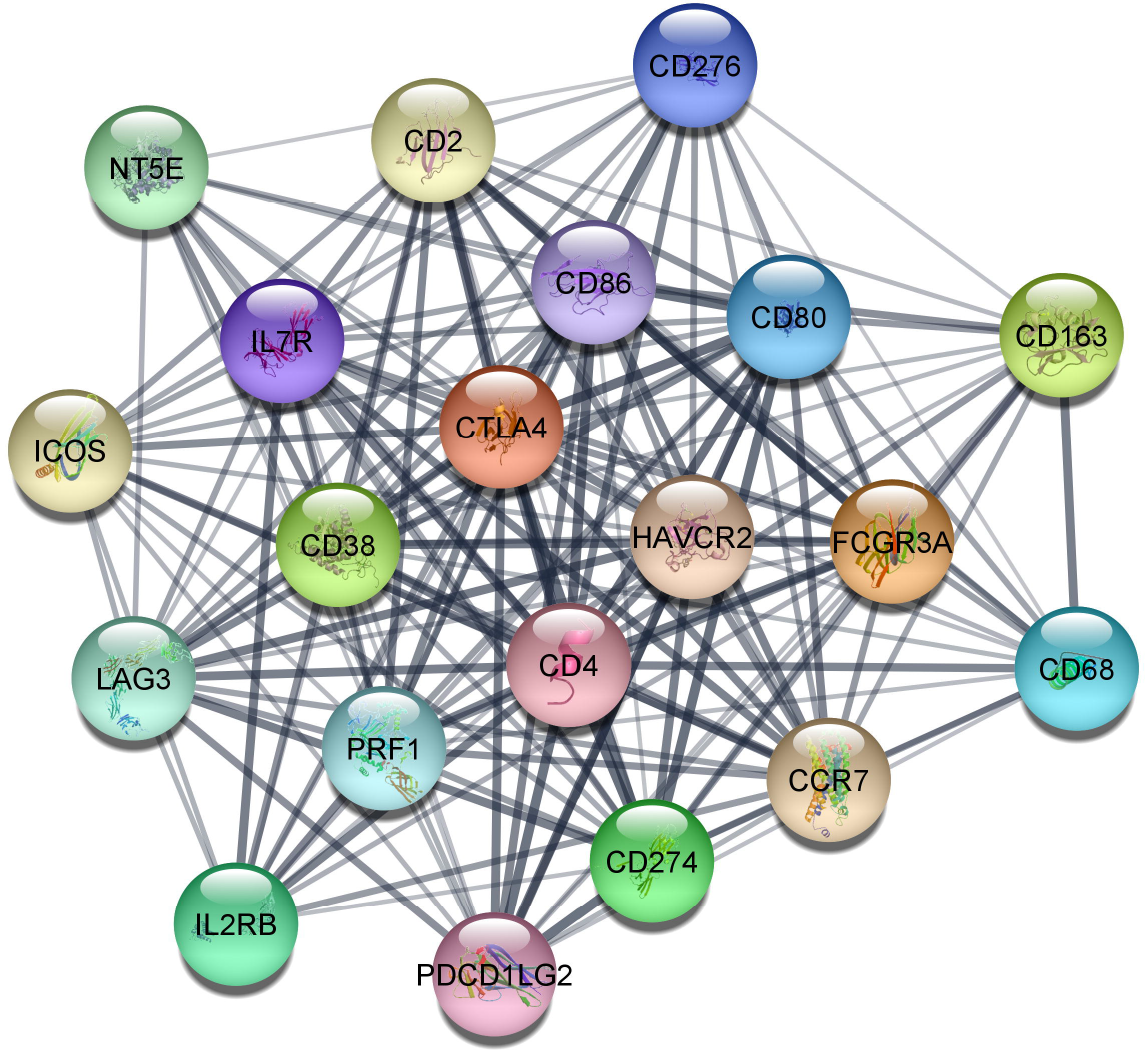

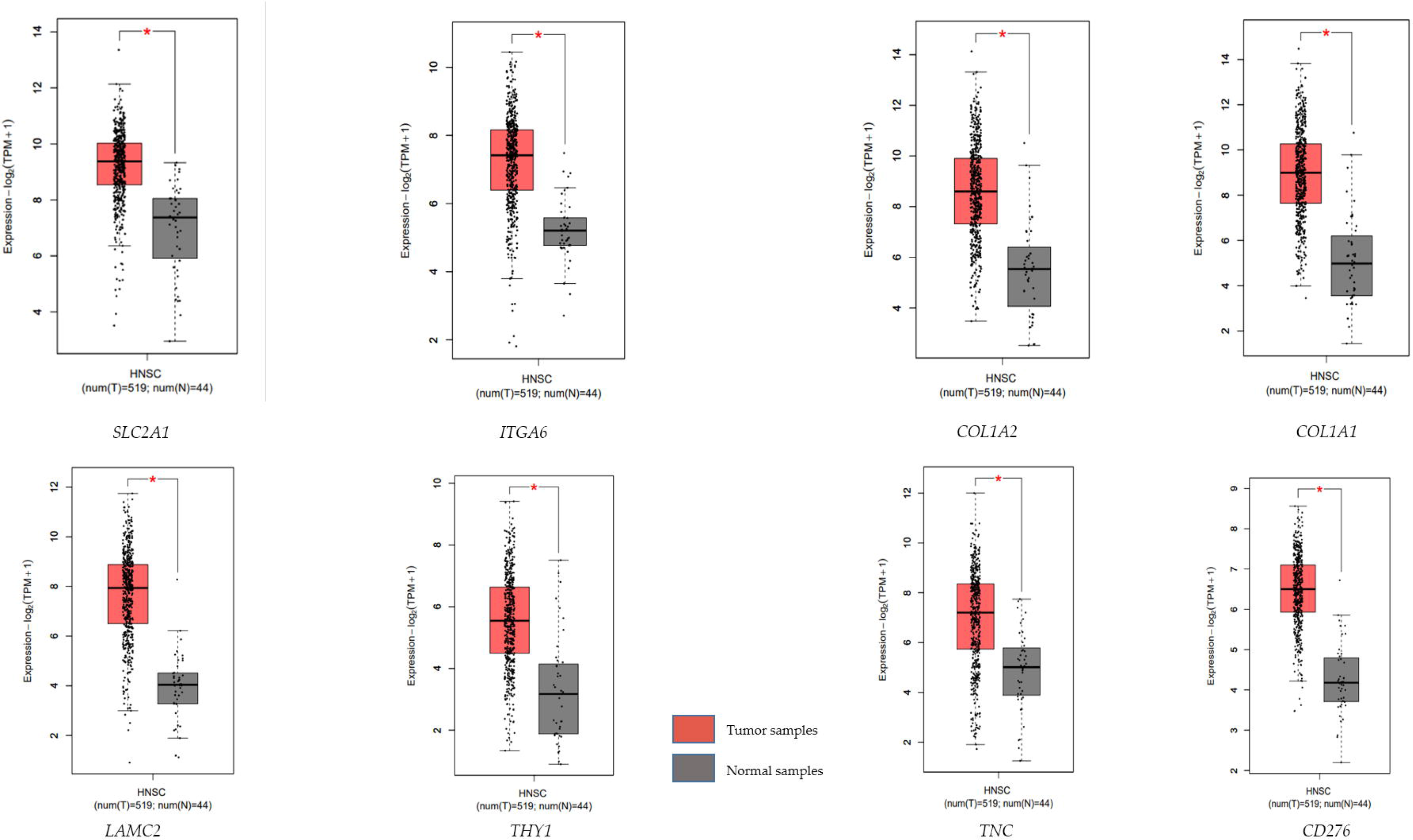

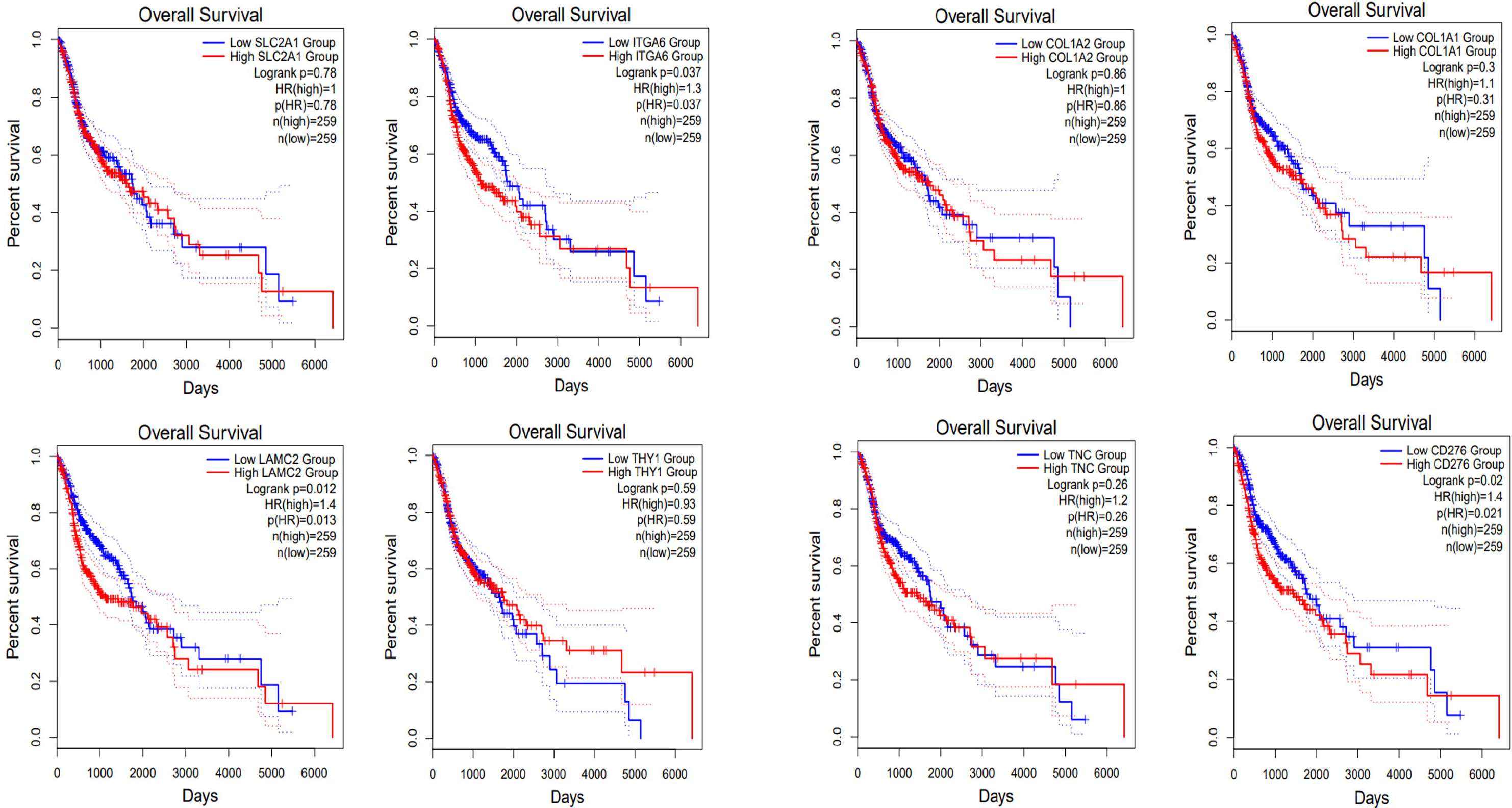

